# Anoxia-Induced VEGF-Release from Rat Cardiomyocites Promotes Vascular Differentiation of Human Mesenchymal Stem Cells

**DOI:** 10.1101/110908

**Authors:** R. Gutierrez-Lanza, V.M. Campa

## Abstract

**Background:** Albeit several studies show that cellular therapy with bone marrow mesenchymal stem cells (BM-hMSCs) improves cardiac function after myocardial infarction (MI), the underlying mechanism is subject of controversy. Here we hypothesized that soluble factors, including VEGF, secreted by cardiomyocytes and BM-hMSCs under a low oxygen environment promote vascular differentiation of BM-hMSCs.

**Methods:** Human BM-MSCs were isolated and expanded *in vitro* by the same procedure we employed to prepare cells suitable for cardiac cell therapy. BM-hMSCs were characterized by flow cytometry and functional analysis. Vascular differentiation was induced by VEGF or by conditioned media from neonate rat ventricular cardiomyocytes (NRVCs) cultured in anoxia and confirmed by immunostaining, tube formation over matrigel and cell migration across transwells. Presence of VEGF in conditioned media was determined by ELISA and activation of VEGF signaling by Western blot.

**Results:** BM-hMSCs used in this study met the criteria recommended by International Society for Cellular Therapy for defining mesenchymal stromal cells. Von Willebrand factor (vWF) expression and tube formation in matrigel indicate that these cells had the capacity to differentiate into the endothelial lineage, which was further enhanced by VEGF and conditioned media from NRVCs cultured in anoxia. Furthermore, condition media and VEGF stimulated cell migration across transwells, which demonstrates the migratory capacity of BM-hMSCs. Finally, when VEGF signaling was blocked by neutralizing anti-VEGF, vascular differentiation of BM-hMSCs was reduced to basal levels.

**Conclusions:** Soluble VEGF released in the culture media after exposure to low oxygen conditions is responsible for endothelial differentiation of BM-hMSCs.

## Introduction

Myocardial Infarction (MI) associated with coronary atherosclerosis remains a leading cause of mortality and morbidity in developed countries. Although, trombolytic therapies have helped to reduce mortality caused by MI, the inability of the heart muscle to regenerate after injury (such as MI), produces a work overload in the heart that decreases enormously the live quality of these patients and frequently leads to congestive heart failure unless the heart is replaced. Today, traditional drug-based therapies only try to reduce this overload by reducing blood pressure or increasing contractibility, but do not facilitate healing of the injured heart and therefore only delay the onset of the final heart failure. Recently, cell-based therapies with stem cells have appeared as an alternative to heart transplatation for their potential to regenerate the damage heart and bone marrow mesenchymal stem cells (BM-hMSCS) have been proposed as a viable source of cells.

Mesenchymal Stem Cells are non-hematopoyetic cells that represent approximately 0.1% of bone marrow mononuclear fraction and have the ability to differentiate into different lineages including bone [1], cartilage, fat, tendon, [2] and myocardium [3, 4]. These cells also have several characteristics that make them very attractive for cardiac cell therapy: they can be easily isolated and expanded *in vitro* up to 40 doublings [5], they can be used as an allograft and they can differentiate to cardiomyocytes and endothelial cells [3, 6].

Several pre-clinical and clinical trials using BM-MSCs show an improvement in several cardiac functions such as ejection fraction, contractibility and end systolic and diastolic volumes [7-13]. However, the extremely low efficiency and weak evidence seen for *in situ* differentiation of BM-MSCs to cardiomyocytes (review by Psaltis et al. [14]), which in any case is sufficient to explain an improvement in cardiac functions, have made many researchers to rethink the mechanism that account for the functional benefits. Thus, it has been proposed that BM-MSCs may protect cardiomyocytes from apoptosis [15], stimulate the regenerative capacity of resident stem cells [16], modulate the inflammatory response associate with MI [17, 18], modulate a remodeling of the scar [19, 20] and contribute to restore the capillary network damage during the MI [20-22].

Here, we have used an *in vitro* model of MI to demonstrate that when deprived of O_2_, cardiomyocytes secrete angiogenic factors, such as VEGF, that stimulate the capacity of BM-hMSCs to differentiate into the endothelial lineage. This suggests that re-vacularization is, at least in part, responsible for the benefit obtained with these cells and therefore future efforts in designing therapies should focus in enhancing and exploding this unique capacity of BM-MSCs.

We also studied the capacity of BM-hMSCs to differentiate into cardiomyocytes when cultured with bona fide cardiomyocytes or to protect them but we did not find evidences supporting any of these mechanisms.

### Material and Methods

#### Isolation and Culture of Mesenchymal Stem Cells

Bone marrow samples were collected at Hospital Rio Ortega (Valladolid) from ileac crest of donors following standard procedures after obtaining informed consent according to the Center Ethic Committee. Isolation and culture of BM-hMSCs was performed as described previously [23]. Briefly, mononuclear cells were purified from bone marrow samples by a gradient discontinuous density centrifugation in Ficoll (d=1.0.73 g/ml). Then, mononuclear fraction at the interphase was recovered and cultured with mesenchymal media [high glucose Dulbecco's Modified Eagle Medium (4.5 g/l glucose, Gibco) supplemented with 20% FBS, 2 mM Glutamax™, non essential amino acids x1 (Gibco) and penecilin (100 U/ml) streptomycin (100 μg/ml)] at 37°C in 21% O_2_ and 5% CO_2_ in tissue culture grade plastic (Corning). After two days in culture, BM-hMSCs adhered to the bottom of culture plates and non-adherent cells were rinsed with PBS×l before the media was changed. Thereafter, medium was changed every two days and cells passed by trypsination when they reached approximately 80% of confluence.

For osteogenic differentiation, cells were plated at a density of 5000 cell/cm^2^ and incubated for 10-12 days in mesenchymal media supplemented with 10^−4^mM dexamethasone, 0.2 mM ascorbic acid and 10 mM β–glycerophosphate (all Sigma). The medium was replaced every other day. Alkaline phosphatase activity in differentiated cells was determined by using the Fast Red Substrate Pack (BioGenex) and calcium depositions by the Von Kossa Staining Kit (Bio-Optica).

For adipogenic differentiation, cells were incubated for 48 hours in mesenchymal media supplemented with 1x Insulin-Tranferrin-Selenium (Sigma), 10^−2^ mM caffeine and 10^−3^mM dexamethasone. Then, medium was replaced by mesenchymal media containing 1x Insulin-Transferrin-Selenium and cells were cultured for additional 20 days. The medium was replaced twice a week.

For endothelial differentiation, cells were incubated in low serum cardiac media (see below) supplemented with recombinant human VEGF_165_ (R&D) for 7 days. Alternatively, cells were incubated with conditioned media from NRVCs (see below) for the same period of time. The media was changed every two days.

#### Isolation and Culture of Neonate Myocytes

NRVCs were isolated from Sprague-Dawley rats as previously described [24]. Briefly, ventricles were dissected from 1-2 days old rats, digested 5 times, 15 minutes each, with Collagense A (0.45 mg/ml) and Trypsin (1 mg/ml) in HBSS×1. Cells were pooled, pre-plated for 60 minutes on a non-coated dish to remove fibroblasts, and plated on 1% gelatin coated cell culture plastic dishes in high serum cardiac media [DMEM:F12 (1:1) (Gibco), 0.2% BSA, 3mM Na-pyruvate, 0.1 mM ascorbic acid, 2 mM L-Glutamax^TM^ and penicillin (100 U/ml) streptomycin (100 μg/ml)] supplemented with 10% horse serum and 5% FBS. After 24 hours media was changed to low serum medium (same but with 0.5% FBS) and cell were cultured at 37°C in 21% O_2_ and 5% CO_2_ for additional 24-48 hours before exposure to anoxia. The purity of cultures was routinely determined by immunofluorescence staining with anti-αActinin monoclonal antibody (Sigma) and Alexa488-congugated anti-mouse immunoglobulin secondary antibody (Invitrogen) and only cultures with >80% of cardiomyocytes were exposed to anoxia. To mimic the ischemic injury *in vitro,* cells were exposed to anoxia using a Gaspak Pouch (Becton Dickinson), which catalytically reduced the percent of O_2_ to less than 0.7% in 150 minutes, after the media was changed with fresh medium (4.5 g/l glucose or glucose-free). After exposure to anoxia, conditioned media was recovered and filtered with a 0.22 μmØ low protein binding filter (Millipore), diluted 1:1 in fresh media and store at 4°C for no more than two weeks. Glucose concentration in this samples was determined using a glucose sensor BREEZE^™^ (Asensia^®^) following manufacturer′ s instructions.

#### Flow Cytometry

Cells were trypsinized, washed with PBSx1, resuspended in 0.5 ml of PBSx1+ 0.5% BSA (10^6^ cells/ml) and incubated with Fluorescein-5-isothiocyanate– (FITC) or phycoerythrin- (PE) conjugated antibodies against indicated clusters of differentiation (Table 1). Cells were analyzed with a Coulter^®^ Epics^®^ XL^™^ flow cytometer and fluorescence intensity of samples compared with the intensity of appropriated isotype controls. PE and FITC Mouse IgG1 (Becton Dickinson) were used as isotype controls.

**Table 1.** Antibodies and dilutions used for cytometry.

|  |  |
| --- | --- |
| Anti-CD90 FITC | (1:250, ref: 555595, BD Pharmigen) |
| Anti-CD105 FITC | (1:5, ref: 105F, Immunostep) |
| Anti-CD166 PE | (1:5, ref: 555647, BD Pharmigen) |
| Anti-CD34 PE | (1:5, ref: 345802, BD Pharmigen) |
| Anti-CD45 Cy5 | (1:5, ref: 332784, BD Pharmigen) |
| Anti-HLA-DR PE | (1:20, ref: CYT-DRPE, Cytognos) |
| Anti-CD117 PE | (1:20, ref: 340529, BD Pharmigen) |
| Anti-CD133 PE | (1:5, ref: 12-1338-71, eBioscience) |
| Anti-CD31 PE | (1:5, ref: 340297, BD Pharmigen) |
| Anti-CD106 PE | (1:5, 555647, BD Pharmigen) |
| IgG1 Isotype Control PE | (BD Pharmigen) |
| IgG1 Isotype Control FITC | (BD Pharmigen) |

#### Cell Viability

After exposure to anoxia, cells were stained for 15 minutes at room temperature with 4 μM Ethidium Homodimer-2 (EtHD-2, Invitrogen) and 10 μM Hoechst (Invitrogen) in PBSx1. Next, cells were washed with PBSx1, fixed with 4% PFA, and blue and red fluorescence was detected by epi-fluorescence microscopy. While Hoechst is membrane permeable and stains all cell nuclei, EtHD-2 only enters cells with damaged membrane and undergoes a 40-fold enhancement of fluorescence upon binding to nucleic acids. EtHD-2 is excluded by the intact membrane of live cells and therefore only stains dead cells.

Viability of BM-hMSC was analyzed before or after differentiation by staining for 30 minutes at 37°C with 5 μM Chloromethyl Fluorescein Diacetate (CMFDA, Invitrogen) and 10 μM Hoechst (Invitrogen) in serum-free culture media. Next, cells were washed with PBS×1, fixed with 4% PFA, and green fluorescence detected as above. CMFDA is a non-fluorescent molecule permeable to cell membranes, but once inside a live cell it binds to thiol group of reduced glutathione, which retains it inside the cell. In addition, intracellular esterases cleave off the acetates, releasing a green fluorescent product inside live cells.

#### ELISA

Concentration of VEGF in conditioned media was determined with the Quantikine rat VEGF immunoassay (R&D), following manufacturer ′ s instructions. Briefly, 50 μl per well of culture supernatants or VEGF standards were added to an ELISA plate and incubated for 2 hours. Next, the plate was washed extensively, incubated with 100 μl an enzyme-conjugated sandwich antibody for 1 hour, washed again and incubated with 100 μl of substrate until color developed. Finally, reaction was stopped before measuring absorbance at 450 nm.

#### Immunostaining

After cells were fixed in 4% PFA for 15 minutes at room temperature, non-specific binding was blocked for at least 1 hours with blocking buffer (PBS×1, 50 mM glycine, 2% BSA, 2% goat serum, 0.01% Na-azide). Next, cells were incubated overnight at 4°C with primary antibodies [anti-vWF (DAKO), anti-VEGF (Santa Cruz), anti-KDR (Santa Cruz), anti-CD90 (BD Pharmigen), anti-αActinin (SIGMA), anti-Connexin43 (Santa Cruz), anti-N-Cadherin (gift from Dr. Mark Mercola)] diluted 1:100 in washing buffer (blocking buffer diluted 1:10 in PBS×1) and immune complexes were detected with appropriated Alexa488, Alexa594 and Alexa647-congugated secondary antibodies (Invitrogen) diluted 1:250 in washing buffer. Cell nuclei were stained with with 0.5 μg/ml DAPI (4′,6′- diamidino-2-phenylindole).

#### In vitro angiogenesis

The angiogenic capacity of BM-hMSCs was evaluated with the CHEMICON^®^ *In Vitro* Angiogenic Assay Kit according to manufacturer′ s instructions. Briefly, 50 μl of cold ECMMatrix^™^ were transferred to each well of a 96 well tissue culture plate and incubated at 37°C for 1 hour to allow the matrix solution to solidify. Next, 5000 BM–hMSCs resuspended in 150 μl of conditioned or VEGF-supplemented cardiac media were seed on top of the layer of gel and cultured until tubular structures developed; usually between 4 to 6 hours. Finally, cells were fixed with 4% PFA and formation of tubular structures analyzed by light microscopy. As recommended by the manufacturer, the branch points formed were counted as a way to quantitate the progression of angiogenesis.

#### Cell migration

The migratory property of BM-hMSCs was analyzed by using the BioCoat^TM^ Endothelial Cell Migration System (Becton Dickinson) following manufacturer′ s instructions. To maximize the chemotactic response of BM-hMSCs, cells were starved in serum free media for 5 hours before setting the assay. Then, 20.000 cells were resuspended in 250 μl of fresh serum-free cardiac media and seeded on top of the transwells and next 750 μl of conditioned of VEGF-supplemented medium were added to the plate. Finally, the plate was incubated for 24 hours in a tissue-culture incubator and cells that migrated across the transwells were loaded with CMFDA and detected with epi-fluorescence microscopy.

#### Western blot analysis

Lysates were prepared by incubating cells in TNE x1 with 1mM NaO_3_VO_4_, 5mM NaF, 1mM PMSF and Aproteinine and Laupeptine, on ice for 15 min, cleared by centrifugation at 12.000 rpm in a bench top centrifuge, frozen in dry ice and thawed at 4°C. Protein concentration was determine by Bradford and 30 μg per lane of lysates were resolved in a 10% SDS-PAGE, transferred onto nitrocellulose membranes (Protan^®^;Whatman). Membranes were blocked with 3% BSA for 1 hour at RT and then proved overnight at 4°C with primary antibodies [anti-phospho–Akt Ser473 (Cell Signaling), anti-phospho-Akt Thr308 (Cell signaling), anti-phospho-p38 Tyr182 (Santa Cruz) and anti-p38 (Santa Cruz)] diluted 1:1000 in 3% BSA. Finally, specific bands were detected using HRP-conjugated antibodies (1:10,000 in 5% non-fat powder milk) and the ECL detection system (GE Healthcare). Membranes were stripped by a 20 min incubation with 1M glycine-HCl, pH 2.5 and reprobed again with anti-Akt (Santa Cruz) and anti-p38 antibodies (Santa Cruz) diluted 1:1000 in 3% BSA.

#### Apoptosis

The percentage of dead cells in co-cultures of BM-hMSCs and NRVCs was determined with the PE Annexin V apoptosis detection kit (Becton Dickinson) after BM-hMSCs were labeled with a CD105 FITC-congugated antibody. Briefly, 10^5^ cells were resuspended in 100 μl of 1x binding buffer and 5 μl of PE-AnnexinV and 5 μl of 7-AAD solutions were added. After a 15 minutes incubation, additional 400 μl of 1x binding buffer were added and cells analyzed by flow cytometry with a Coulter^®^ Epics^®^ XL^TM^ flow cytometer. The percentage of Annexin V+ cells was calculated in the CD105- (mesenchymal) and CD105- (cardiac) populations.

#### RT–PCR

Total RNA was extracted from cell cultures with Trizol^®^ reagent (Gibco). One microgram of RNA was converted to cDNA with the QuantiTect^®^ Reverse Transcription Kit (QIAGEN), which also eliminates remaining gDNA in the sample, in a 20 μl reaction following manufacturer′ s instructions. Then, PCR was performed with 2 μl of the cDNA synthesis reaction, 0.2 mM dNTPs, 2.5 U TaqPol (Fermentas), 1.5 mM MgCl_2_ and 0.05 μM of forward and reverse primers in a total volume of 25 μl. Primers sequences (all 5′-3′, forward, reverse), reaction conditions (annealing and elongation temperature and time) and product sizes are as follows: human MHCα: 5′-CAC CAA CCT GTC CAA GTT CC-3′, 5′-GTT GGC AAG AGT GAG GTT CC-3′, 57°C 30′′, 72°C 60′′(174 pb); rat MHCα: 5′-CAC CAA CCT GTC CAA GTT CC-3′, 5′-GCA ACA GCG AGG CTC TTT CTG-3′, 57°C 30′′, 72°C 60′′(174 pb); rat&human GADPH: 5′-GTC GGT GTG AAC GGA TTT G-3′, 5′-ACA AAC ATG GGG GCA TCA G-3′, 58°C 60′′, 72°C 90′′, (397 pb); human KDR: 5′- CCC ACG TTT TCA GAG TTG GT-3′, 5′-TCC AGA ATC CTC TTC CAT GC-3′, 54°C 40′′, 72°C 40 (123 pb). The PCR was performed with a Biometra^®^termocycler. The product size was confirmed by running 10 μl of the sample on a 1% agarose gel electrophoresis.

#### Statistical analysis

All data are presented as mean ± SD. Statistical significance was calculated using a two tailored Student *t* test for comparison between two groups. A probability value of <0.05 was considered significant.

## Results

### Characterization of BM-hMSCs

BM-hMSCs were isolated from bone marrow by centrifugation in a discontinuous density gradient of Ficoll and cultured according to the standard protocol for preparation of BM-hMSCs used for cardiac cell therapy at Hospital Rio Ortega (Valladolid). Bright phase micrographies from cultures in passage P1 show cells with a spindle-shape morphology and a length of approximately 200μM (Figure 1A, B). This fibroblast-like morphology is consistent with the expected morphology of BM-MSCs and was conserved for more than 7 passages (not shown) but, in agreement with our approved protocol for cardiac cell therapy, only cells from P1 to P4 were used in this study. Cells were further characterized according to criteria recommended by the International Society for Cellular Therapy [25]. As expected, these cells showed adherence to plastic (Figure 1A, B). Alkaline phosphatase (AP) expression was induced (Figure 1C, D) and calcium deposits (Figure 1E, F) formed when cells were cultured in osteogenic media for 10-12 days, indicating differentiation to osteoblasts. Cell viability was assayed before and after differentiation and in both cases was higher than 99% (Figure 1G. H), indicating that cells still do not show signs of senescence at passage P4. Lipidic vacuoles accumulation inside cells after adipogenic protocol indicates that cells had also the capacity of differentiate to adipocytes (Figure 1I) and von Willebrand Factor (vWF) staining of cells cultured in cardiac media media shows spontaneous differentiation into the endothelial lineage (Figure 1J). Finally, flow cytometry (Figure 1K) showed that, as espected, cells expressed the surface antigens CD90, CD105 and CD166 (typically expressed by MSCs), and lack the expression of CD34, CD45, HLA-DR, CD117 and CD133 (expressed by hematopoietic and endothelial precursors).

**Figure 1:**
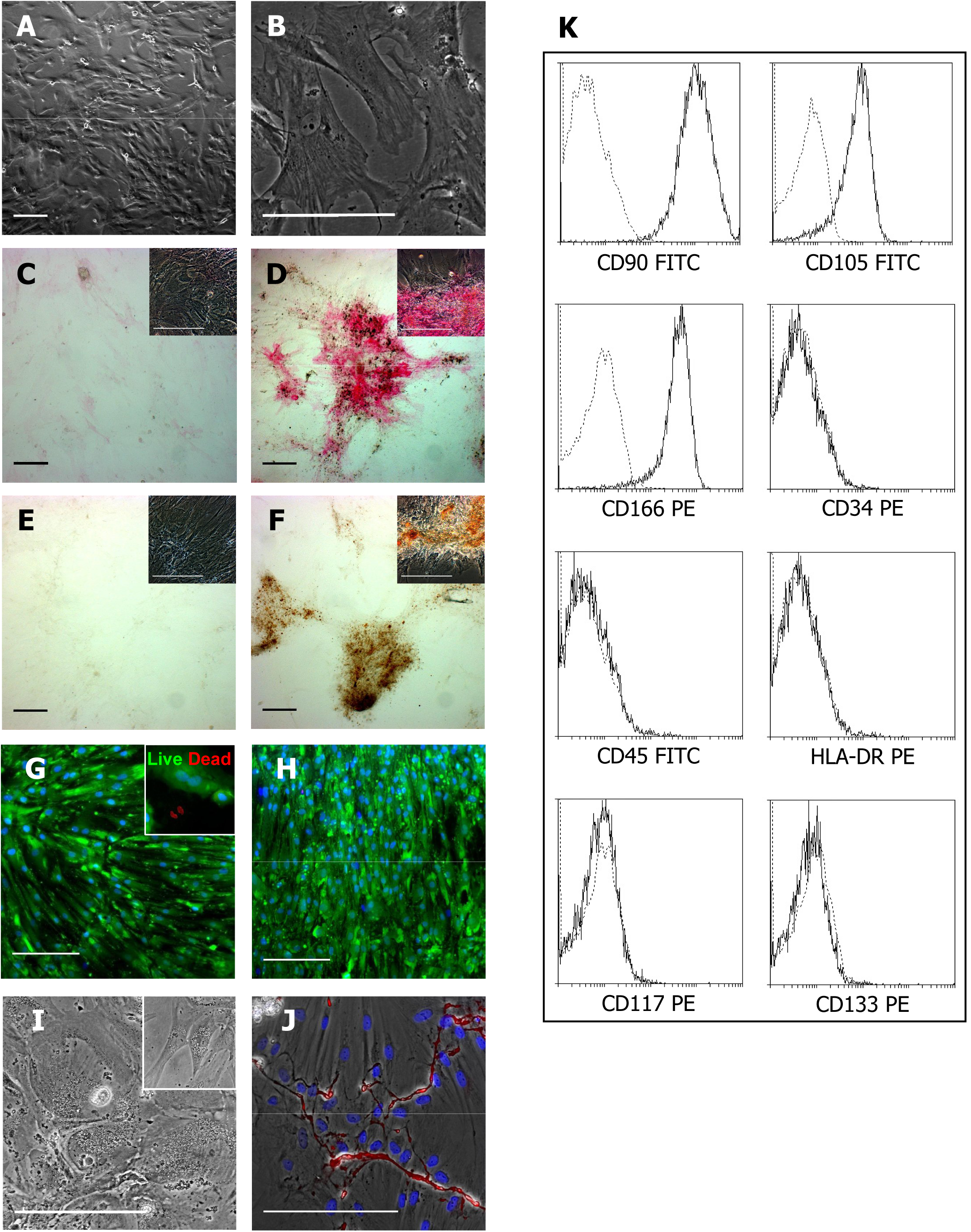
Characterization of BM-hMSCs A-B) Bright phase micrography of P1 BM-hMSCs. **C-H)** BM-hMSCs were incubated for 10 days in osteogenic media (D, F, H) or in control media (C, E, G) and stained with Fast Red for alkaline phosphatase expression (C-D), with Von Kossa for calcium depositions (E-F) or with CMFDA and EtHD for cell viability (G-H). **I)** Bright phase micrography of BM-hMSCs incubated for 14 days with adipogenic media. Insert shows detail of lipidic vacuoles; comprare with 1A **J)** BM-hMSCs were cultured for 10 days in DMEM:F12 supplemented with 10% FBS and then stained with rabbit anti-vWF antibody (red) and DAPI (blue). Bar scales represent 200 μm**K)** Expression of indicated CD markers in undifferentiated hMSCs was analyzed by flow cytometry.

Although flow cytometry analysis indicates that cultures are a fairly homogeneous population we performed a clonogenic analysis to further ensure the stemness of these cells. With 11 clones obtained out of 228 cells plated at a density of 1 cell per well the cloning efficiency was 4.8%.

The differentiation ability of 6 of these clones was analyzed after passing and culturing them for an additional week. Four of them expressed CD90 when cultured in regular media and retained the ability to differentiate to osteoblasts when cultured in osteogenic media or to endothelial cells when cultured in media supplemented with VEGF, whereas the other 2 remaining clones, although still expressed CD90 and vWF, had lost the ability to differentiated to osteoblast and collapsed before reaching confluence (Figure 2).

**Figure 2:**
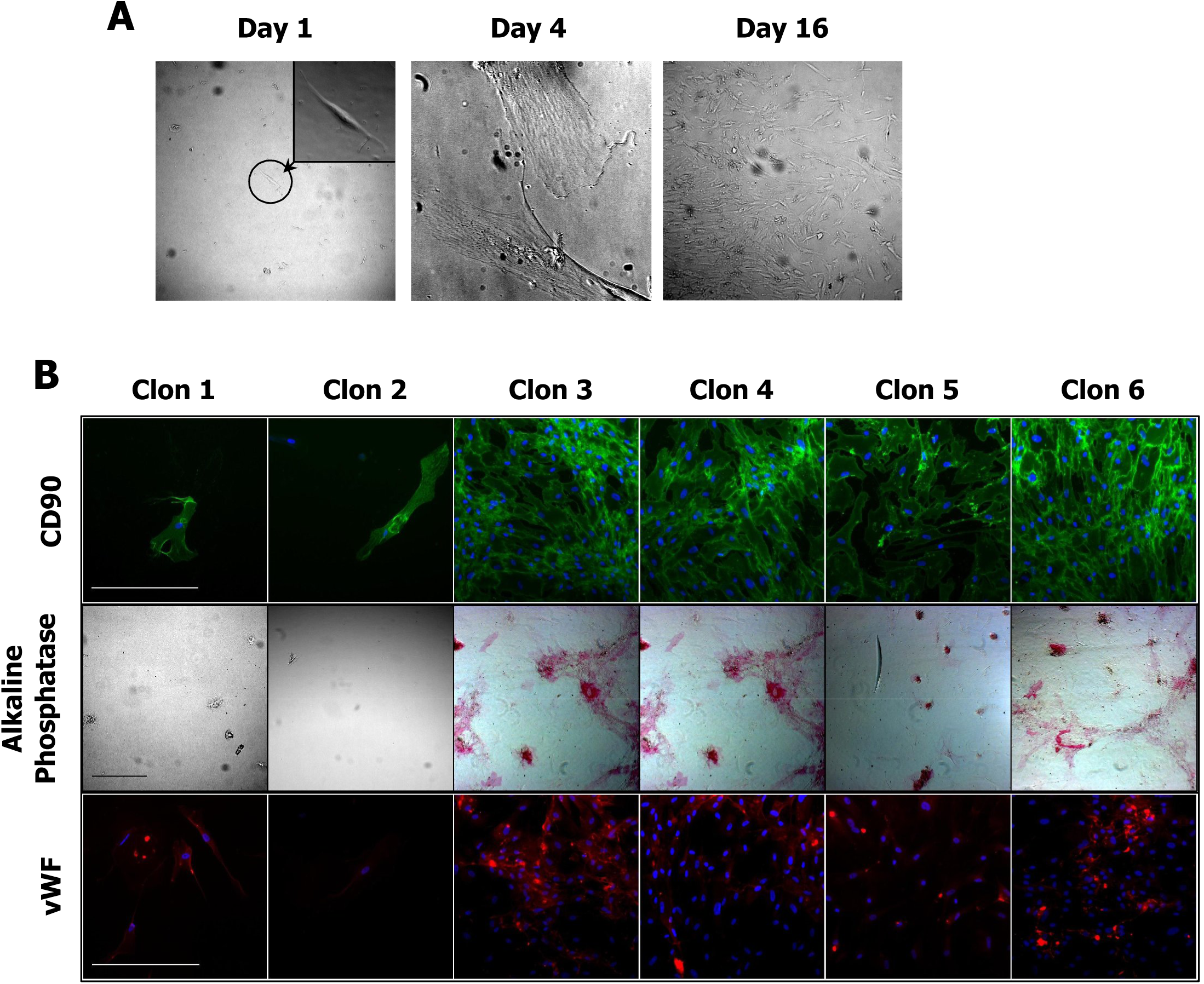
Clonal differentiation of BM-hMSCs A) P1 BM-hMSCs were trypsinized and plated in 96 well tissue culture plates at a density of 10 cells/ml in 100 μl of media. Clonal expansion of single BM-hMSCs was monitored every other day for a period of up to two weeks. **B)** When different clones reached 70-90% of confluence, cells were trypsinized again and plate at a 1 to 3 dilution in a new 96 well plate. Next, each clon was cultured for 7 days in control media, osteogenic media or control media supplemented with VEGF. Finally, cells were fixed and stained with mouse anti-CD90 FITC-congugated antibody (green), Fast Red (for alkanine phosphatase expression) or rabbit anti-vWF antibody (red), as indicated.

Taken together, our results indicate that this is a fairly homogenous population of BM-hMSCs, which has the capacity of self-renewal and the ability to differentiate to various lineages, including the endothelial. However, the collapse of 2 clones and the impossibility of growing indefinitely any of the remaining 4 clones, or the original primary culture, suggests cellular senescence in late passage cultures. Cellular senescence of BM-MSCs after prolonged culture it has been previously reported [5].

### Soluble factors released from NRVCs induce endothelial differentiation of BM-MSCs

In order to study the bases of the therapeutical benefit seen after cardiac cell therapy with BM-hMSCs we used an *in vitro* model of MI [26] developed with ventricular cardiomyocytes isolated from neonate rats (NRVCs). As expected, with more than 80% positive cells, NRVCs stained for α-Actinin, and adjacent cells showed Connexin43 and N-Cadherin (Figure 3A) and beat synchronously (not shown).

**Figure 3:**
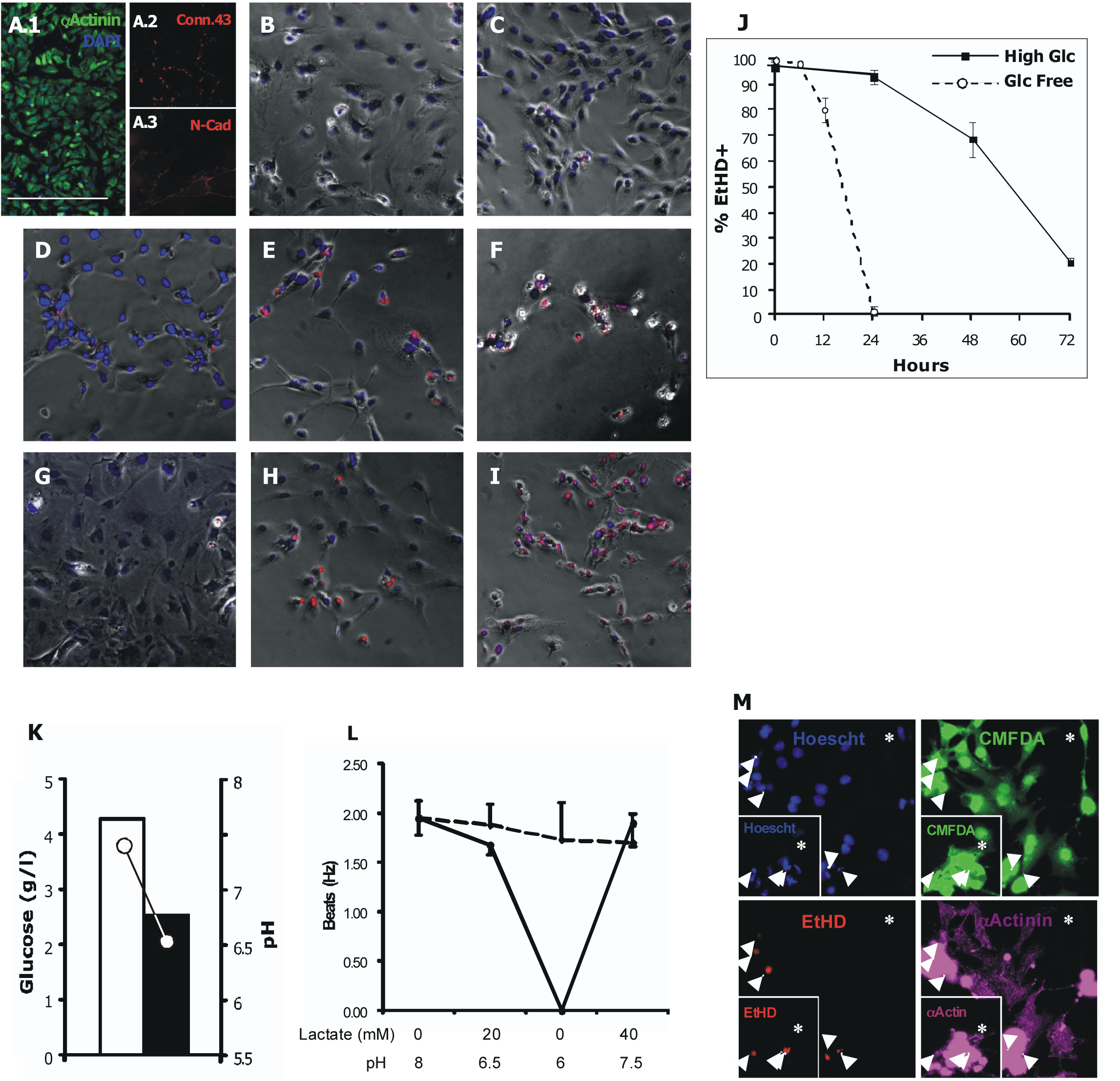
Cell death of NRVCS after exposure to anoxia. **A)** 72 hours after isolation NRVCs were stained with mouse anti-αActinin antibody (green) and DAPI (blue) (A1), mouse anti-Connexin43 antibody (red) (A2) and rabbit anti-N-Cadherin antibody (red) (A3). **B-J)**NRVCs
were incubated in glucose free DMEM (B, D-F) or in high glucose DMEM (4.5 gr glucose/L) (C, GI) under normoxic (O_2_ ≈20%) (B-C) or anoxic conditions (O_2_ 20%) (B-C) or anoxic conditions (O_2_ <1%) (D-I) for 6 (D), 12 (E) 24 (B, G) or 48 (H) or 72 (C, I) hours and then dead cells were stained with EtHD. EtHD positive cells were counted and the percentage of cell death calculated (J). **K)** Glucose concentration and pH in or conditioned media from NRVCs culture for 24 hours in normoxia or anoxia in high glucose DMEM. L) Lactate concentration (dashed line) and pH (filled line) was sequencially altered as indicated 15 min before frequency of contraction was determined. M) NRVCs were incubated in cardiac media for 36 hours under anoxic conditions and then live cells were stained with CMFDA (green), dead with EtHD (red), nuclei with Hoechst (blue) and cardiomyocytes with mouse anti-αActinin antibody (far-red, shown as magenta).

When NRVCs were cultured in anoxia, the number of EtHD positive cells, which was used as cell death marker, increased with time after exposure to anoxia. When cells were exposed to anoxia in media containing glucose (4.5 g/l), 6.8%±2.8% of NRVCs stained with EtHD after 24 hours, 31.3%±6.8% after 48 hours and 79% ±1.5% after 72 hours. In contrast, when cultured in normoxia, the percentage of EtHD positive cells after 72 hours was only 3.3% ±1.8% (Figure 3B, D-F, J). In addition, levels of glucose in the media and pH decreased after exposure to anoxia but not in cells kept in normoxia (Figure 3K). During anoxia cells stop beating, probably because lactate accumulation, and resumed beating when normal O_2_ level were restored. In fact, NRVCs stopped beating when cultured in acidified fresh media and resume beating when physiological pH was restored (Figure 3L). Interestingly, presence of lactate in the media did not decrease the frequency of contraction, suggesting that media acidification it is the main responsible of the inhibition of contractility. When cultured in glucose free media, the percentage of EtHD positive NRVCs increased much faster after exposure to anoxia, with 1.9%±0.9% of positive cells after 6 hours, 20% ±4.8% after 12 hours, and 98.4% ±1.7% after 24 hours (Figure 3C, G-I, J). During normoxia only 0.46% ±0.15 of cells stained for EtHD (Figure 3C). This results support the idea that, when expose to low O_2_ conditions, cardiomyocytes depend completely on glycolysis to obtain their energy and enter in a lethargic state, which may reflect the inert area of the peri-infarcted region, that might end with cell death when energy production is not enough to meet the minimal metabolic needs, or resume contractility if O_2_ levels recover and aerobic metabolism restores. Logically, a therapeutic benefit involving these cells only can be obtained before cell death is irreversible. Therefore, the remain experiments of this study were performed during the few hours after cell death started to be detected: 36 to 48 hours in media containing glucose and 6 to 12 hours in glucose free media; a period of time we hypothesized relates to the therapeutic window.

As 10 to 20% of cells did not stain for α-Actinin, we wanted to be sure that cardiomyocytes were dying during the period of time mentioned above. When live cells were stained with CMFDA, dead cells with EtHD and cardiomyocytes with α-Actinin, 36 hours after exposure to anoxia in glucose containing media, we observed that the vast majority of dead cells were cardiomyocytes (Figure 3M, indicated by arrows), whereas than less than 1% of α-Actinin negative cells stained for EtHD. The reduction of the percentage of α-Actinin positive cells after exposure to anoxia further confirmed this idea (not shown).

In response to low O_2_ levels, cells respond with stabilization and activation of transcription factor HIF1α. Once activated, HIF1α translocates to the nucleus where it binds to hypoxia response elements (HRE) and induces transcriptional activity of genes involved in response to hypoxia, including VEGF [27]. Interestingly, VEGF signaling is indispensable for many aspects of vasculogenesis and angiogenesis, including endothelial differentiation and tube formation [28] and it has been shown to promote differentiation of BM-hMSCs into endothelial cells [6]. Therefore, to check if NRVCs respond to low O_2_ by secreting soluble factors, as VEGF, that can promote vascular differentiation of BM-hMSCs, we analyzed VEGF production in NRVCs after exposure to anoxia. Levels of VEGF in culture supernatants increased from 85ng/ml, when cells were cultured in normoxia, to 1013ng/ml after exposure to anoxia for 24 hours (Figure 4A). This is a more than 10 fold induction. Furthermore, NRVCs secreted 949ng/ml of VEGF during the 24 hours following exposure to anoxia, indicating that VEGF-expression persists for a long period after being stimulated by exposure to low O_2_ levels. Immunofluorescence staining against VEGF further confirmed that VEGF is expressed by NRVCs after exposure to anoxia (Fgure 4B-C). Costaining with vital dye CMFDA demonstrates that fluorescence signal arise from live cells and is not due to autofluorescence of dying cells under anoxic conditions.

**Figure 4:**
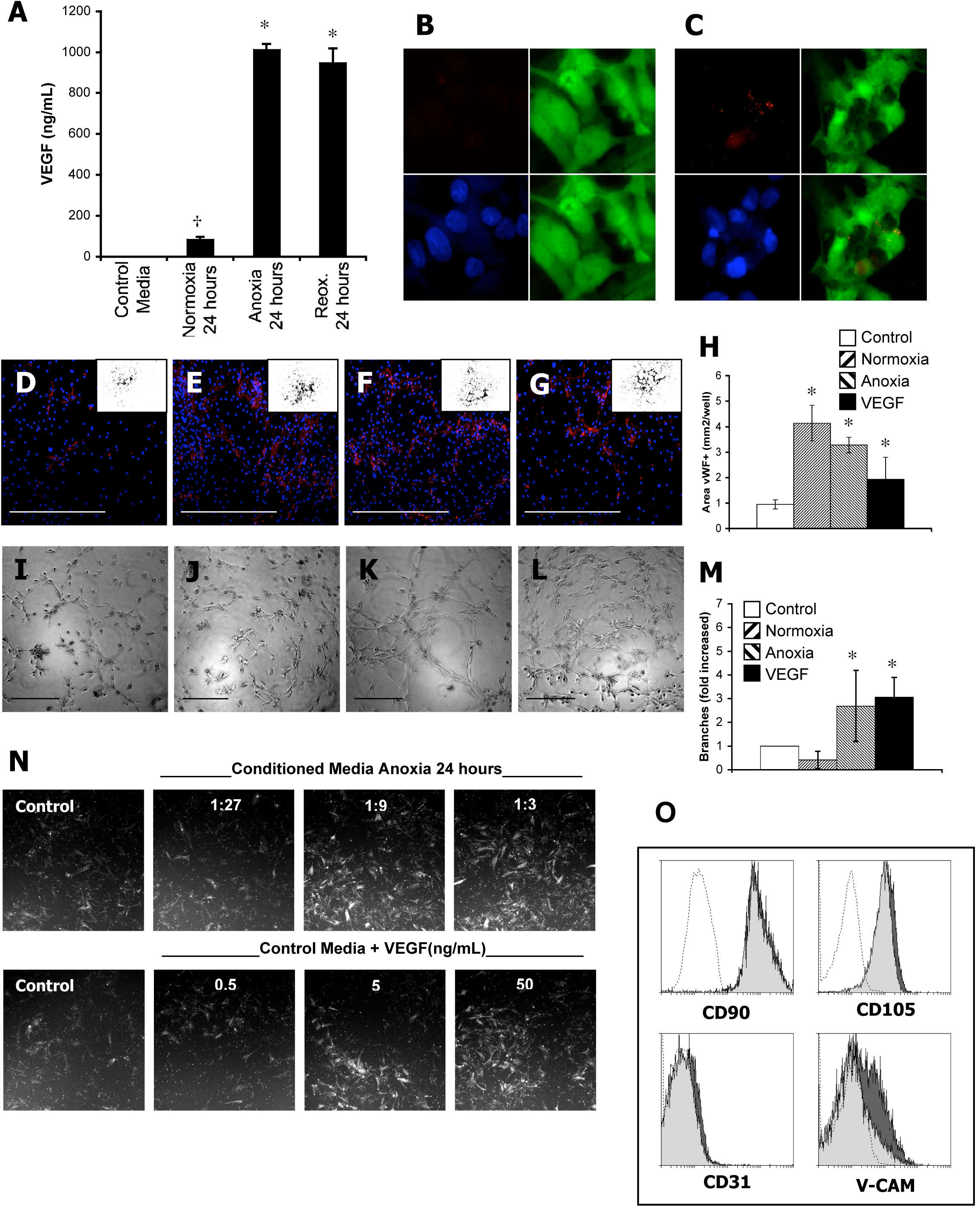
Vascular differentiation of BM-hMSCs A) NRVCs were incubated for 24 hours in Anoxia or in Normoxia, with or without a previous 24 hours incubation in Anoxia (as indicated). Then, soluble VEGF was determined in supernatants by ELISA. ^*^ p<0.001 vs.normoxia; †p<0.005 vs. control media.**B-C)** NRVCs were incubated for 24 hours in Normoxia (B) or in Anoxia (C) and then cells were stained with rabbit anti-VEGF antibody (red), CMFDA (green) and Hoechst (blue). **D-H)** BM-hMSCs were incubated for 7 days in control media (D), or conditioned media from NRVCs incubated for 24 hours in normoxia (E) or in anoxia (F), or in control media supplemented with VEGF (100 ng/ml) (G). Then cultures were fixed with 4% PFA and stained with rabbit anti-vWF antibody (red) and DAPI (blue). Scale bars represent 400 μm. Finally, positive area for vWF was determined in low magnification images of cultures after applying a mask (inserts) in images (H). ^*^ p<0.05 vs. control media.**I-M)** Light microscopy images of tube formation by BM-hMSCs plated at low density (5000 cells/well in a 96 well plate) on top of polymerized ECMatrix^™^ in control media (I) or in conditioned media from NRVCs incubated for 24 hours in normoxia (J) or in anoxia (K), or in control media supplemented with VEGF (100 ng/ml) (L) and cultured for 6 hours. Scale bars represent 400 μm. The capillary tube branch points formed were counted and fold increase calculated (M). ^*^ p<0.05 vs. control media.**N)** Cellmigration capacity of BM-hMSCs in presence of indicated dilutions of conditioned media from NRVCs cultured in anoxia for 24 hours, or indicated concentration of VEGF, across BIOCOAT^™^transwells (3 μM pore). Note that conditioned media and VEGF stimulate the migration in a dose dependent fashion. **O)** Expression of indicated CD markers in BM-hMSCscutured for 7 days in control media (light grey) or in media supplemented with VEGF (100 ng/ml) (drak grey) was analyzed by flow cytometry.

Next, we checked if conditioned media from NRVCs induces vascular differentiation of BM-hMSCs. When BM-hMSCs were cultured during 7 days in conditioned media from NRVCs cultured in normoxia or anoxia, the vWF positive area of cell cultures was 4.14 ±0.31 and 3.28 ±0.86 mm^2^/well respectively (Figure 4E-F, H). In contrast, when BM-hMSCs were cultured for this period of time in fresh cardiac media vWF positive area was only 0.96 ±0.18 mm^2^/well (Figure 4D, H). This is a more than 3 fold increase in each case. As expected, VEGF, with a calculated value of 1.95 ±0.70 mm^2^/well, also increase the vWF positive area (Figure 4G, H).

We also analyzed the angiogenic potential of BM-hMSCs by measuring the capacity to form a capillary network when cultured on matrigel. The number of branches formed by undifferentiated BM-hMSCs in presence of conditioned media from NRVCs cultured in anoxia for 24 hours increased 2.69 ±1.57 times when compared to tubes formed in fresh cardiac media (Figure 4I, K, M). In this case, conditioned media from NRVCs cultured in normoxia (0.42 ±0.36 fold increase) did not have an effect (Figure 4J). Finally, VEGF, as expected, increased the number of branches 3.05 ±0.84 fold (Figure 4L).

In addition, we analyzed the capacity of BM-hMSCs to migrate across a microporous (3.0 μm pore size) fibronectin coated membrane. Cell migration is an essential during both angiogenesis and vasculogenesis, and it is normally stimulated by VEGF in angioblast and endothelial precursors. We observed that BM-hMSCs spontaneously migrate across the membrane and the number of cell that pass through the porous membrane increased in a dose dependent fashion when VEGF or conditioned media from NRVCs cultured in anoxia was used as chemoattractant (Figure 4N).

Finally, we analyzed the expression of mesenchymal markers and endothelial markers in undifferentiated and in VEGF-stimulated to confirm the potential of BM-hMSCs to differentiate into the endothelial lineage. Expression of mesenchymal markers CD90, CD105 was homogeneous in cell cultures and presence of VEGF in media for 7 days did not alter levels of expression of these markers (Figure 4O). Expression of early endothelial marker V-CAM was detected at low level in a small percentage of undifferentiated, which probably represents the spontaneous differentiation of these cell mentioned above, and although levels of expression did not change, the percentage of V-CAM expressing cells increased upon VEGF stimulation, confirming that VEGF stimulates the endothelial differentiation of these cells. The potential for endothelial differentiation of individual cells was confirmed by a clonogenic analysis (Figure 2). On the other hand, expression of late endothelial marker CD31 was not detected nor in undifferentiated neither in VEGF-stimulated cells (Figure 4O), suggesting that terminal differentiation is not complete, at least in cell culture.

Next, we study expression of KDR and VEGF signal in BM-hMSCs. In agreement with data published by Oswald et al. we did not detect KDR expression by flow cytometry (not shown). However, IF staining with anti-KDR antibody showed a weak but specific staining localized in cell membranes (Figure 5A). Furthermore, VEGF-treated but not un-stimulated cells stained for VEGF after protein fixation (Figure 5B-C), indicating that VEGF binds to BM-MSCs. In addition, KDR mRNA was detected in BM-hMSCs (Figure 5D). Finally, we tested the phosphorylation status of Akt and p38. Akt and p38 are downstream elements of VEGF signaling pathway that become activated by phosphorylation in presence of VEGF. As expected, VEGF induced a transient phosphorylation of Akt and p38 that peaked at 30 minutes and returned to basal levels after 1-2 hours (Figure 5E). Conditioned media from NRVCs exposed to anoxia also induced a transient phosphorylation of Akt and p38 that was abolished when conditioned media was pretreated with anti-VEGF antibodies (Figure 5E). Taken together, these results indicate that KDR is being expressed, probably a low levels, in BM-hMSCs and that VEGF signaling pathway is activated by soluble VEGF present in conditioned media from NRVCs cultured in anoxia.

**Figure 5:**
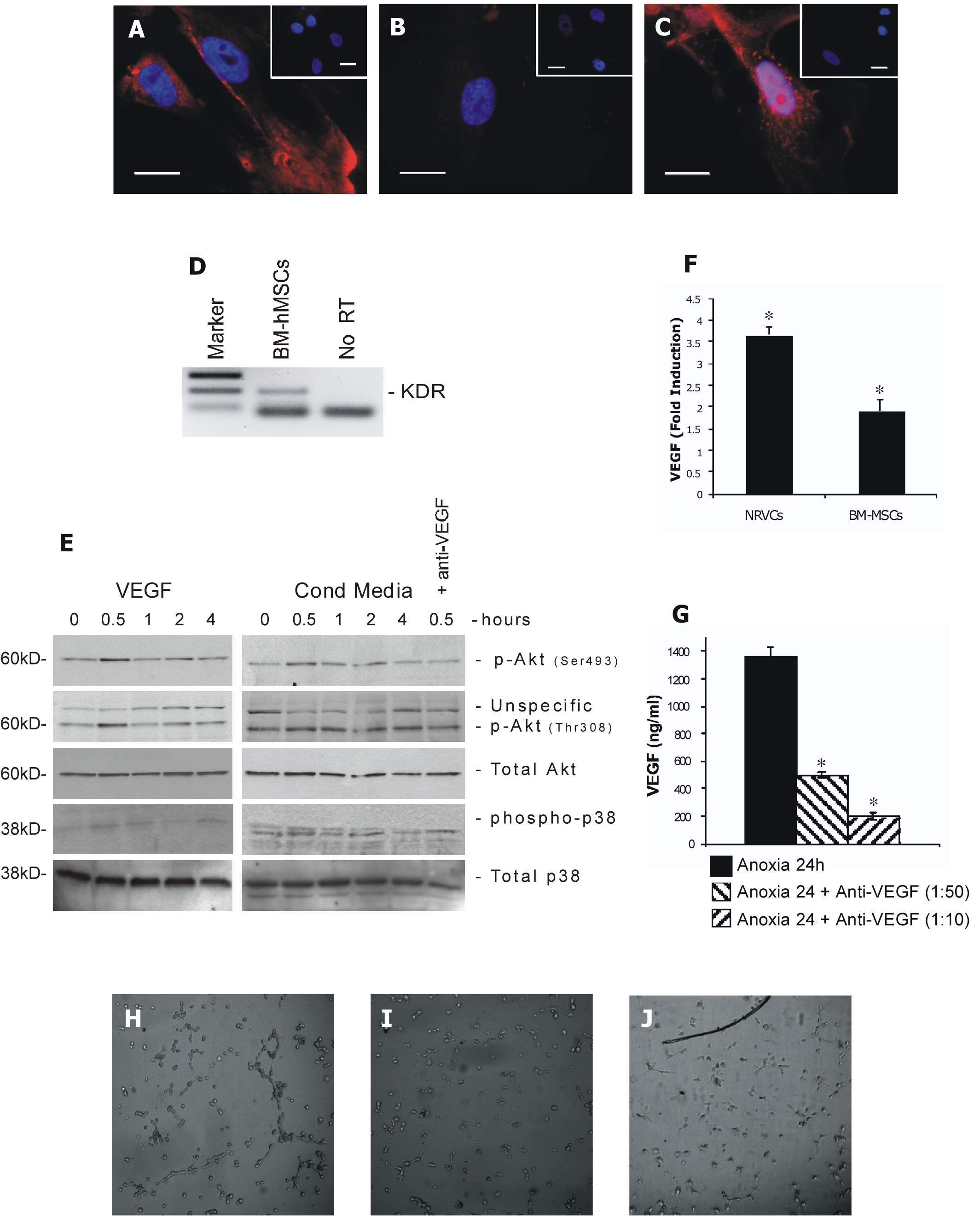
A-C) BM-MSCs were incubated for 1 hour with fresh media in absence (A-B) or presence (C) of VEGF (50 ng/ml). Next cells were fixed with 4% PFA and stained with anti-KDR (B-C) or anti-VEGF antibodies (red) and DAPI (blue). Scale bars represent 25 μm. **D)** Total mRNA was isolated from BM-hMSCs and KDR mRNA was determined by RT-PCR. **E)** BM-hMSCs were stimulated with VEGF (100 ng/ml) or with conditioned media from NRVCs cultured in anoxia in absence or, if indicated, presence of anti-VEGF antibody (1:25) for 24 hours during indicated time and then phosphorylation status of Akt and p38 was analyzed by Western blot. **F)** NRVCs and BM-hMSCs were incubated for 24 hours in anoxia or normoxia. Then, soluble VEGF was determined in supernatants by ELISA. ^*^ p<0.001 vs. normoxia.**G)** Conditioned media from NRVCs cultured in anoxia for 24 hours was incubated with indicated dilutions of anti-VEGF antibody. Then, the antibody was immunoprecipitated before remaining VEGF were determined by ELISA. Note a dose dependent clearing of VEGF. ^*^p<0.001 vs. anoxia conditioned media. **HI)** Tube formation over matrigel of BM-hMSCs plated at low density (5000 cells/well in a 96 well plate) on top of polymerized ECMatrix^™^ in conditioned media from NRVCs cultured in anoxia for 24 hours after blocking soluble VEGF with indicated dilutions of anti-VEGF polyclonal antibody.

Several authors have proposed that BM-MSCs secrete soluble factors that facilitate endogenous repair processes. In agreement with previous reports [29], VEGF expression also was detected in BM-hMSCs and it suffered a twofold increase when cells were exposed to anoxia (Figure 5F). This VEGF not only may act enhancing local angiogenesis but also promoting differentiation of BM-hMSCs in an autocrine way. In fact, when VEGF was removed from conditioned media obtained from NRVCs cultured in anoxia (Figure 5G), the capacity of BM-hMSCs to form tubes on matrigel was completely abolished (Figure 5H-J), indicating that VEGF is the soluble factor responsible of promoting angiogenesis. Taken together, these results show that soluble factors released by NRVCs promote vascular differentiation of BM-hMSCs. They also confirm that VEGF, which is released by NRVCs when exposed to low O_2_ levels, is sufficient to trigger this process, suggesting that it may be the key factor in this process.

Different studies show that BM-hMSCs protect bona fide cardiomyocytes from cell death [15] and that they can differentiate into the cardiac lineage [30]; being both processes dependent on VEGF signaling. Although, we tested if co-culture with BM-MSCs, with conditioned media from BM-hMSCs exposed to anoxia, or with VEGF protects cardiomyocytes by analyzing the number of Annexin V+ cells after exposure to anoxia, we did not find any evidence of protection. The 31.4 ±0.82% of annexin V+ NRVCs after exposure to anoxia for 6 hours did not diminished when NRVCs were co-cultures with BM-hMSCs (41.7 ±1.46%), cultured with conditioned media from BM-hMSCs (35.4%) or stimulated with VEGF (41.2%) ( Supplemental figure 1A-B). Interestingly, BM-hMSCs, with only 7.51 ±0.46% of annexinV+ cells within the CD90+ population, appeared to be more resistant to anoxia that NRVCs. Similar results were obtained when NRVCs were cultured in glucose containing media for 33 hours (not shown). In addition, when we analyzed by RT-PCR the expression of human MHCα, or by IF the expression of α-Actinin, in co-cultures of BM-hMSCs and NRVCs, we did not find any sign of cardiac differentiation in BM-hMSC after one week of co-culture (Supplemental Figure 1C-E).

**Supplemetal figure 1:**
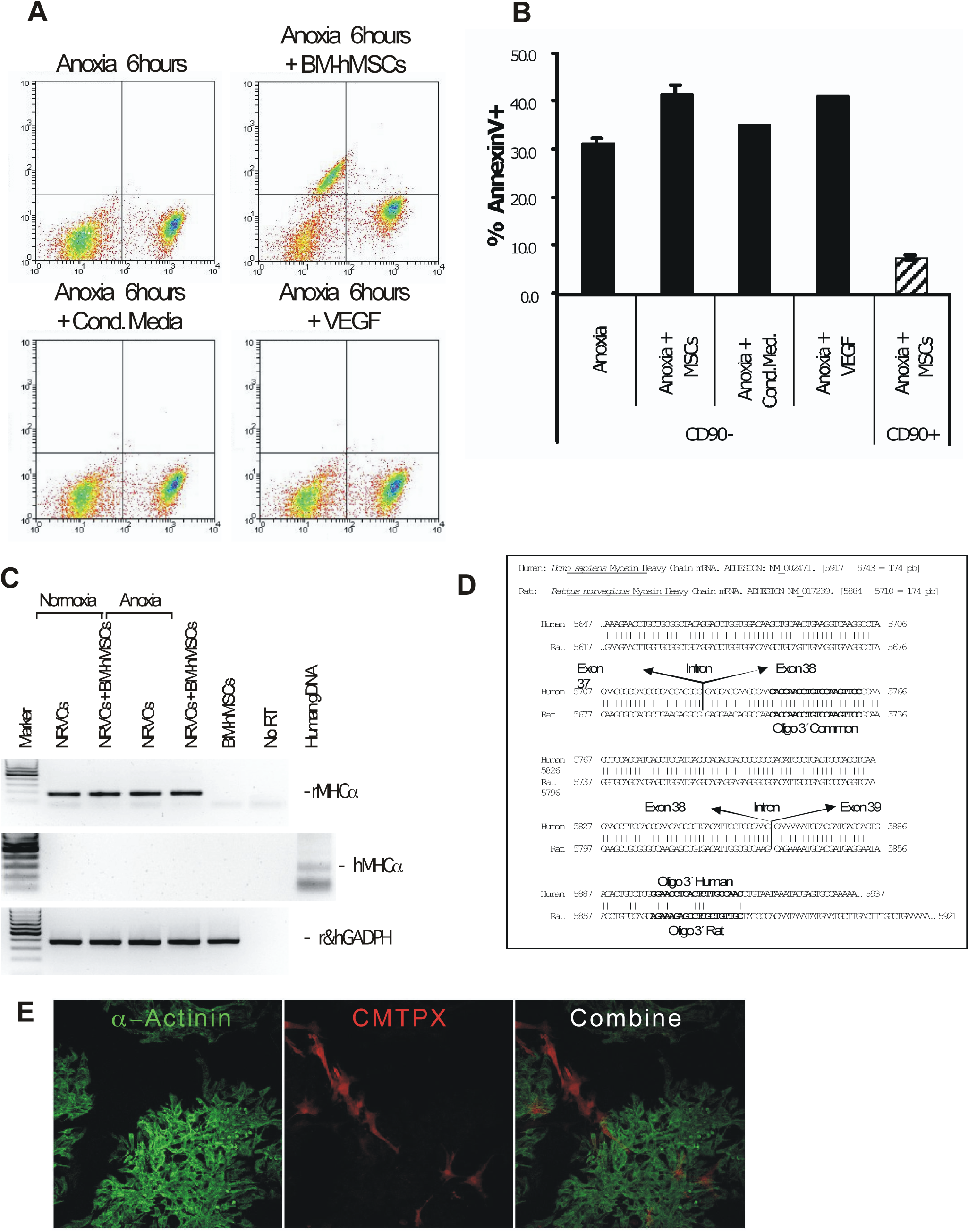

A-B) NRVCs were exposed for 6 hours to anoxia in glucose free media, in co-culture with BM-hMSCs (ratio 10:1), in presence of conditioned media from BM-MSCsculture for 24 hours in anoxia, and in presence of VEGF (100 ng/ml), as indicated. AnnexinV+ cells were identified by flow cytometry and the percentage of positive cells for each condition calculated. **C)** NRVCs were co-culture with BM-hMSCs (ratio 10:1) and expose to anoxia or kept in normoxia for 24 hours. Then, cultured for additional 6 days and rat MHCα, human MHCα, and total GADPH expression were determined by RT-PCR. **D)** Aligned sequences of human and rat MHCαmRNA and primers used for amplification. **E)** BM-hMSCswere loaded with CMTPX and then co-cultured with NRVCs for 5 days. Next, cells were fixed with 4% PFA and stained with anti α-actinin monoclonal antibody (red).

### Discussion

Several clinical trials indicate that cardiac cell therapy with BM-hMSCs after MI improves cardiac function. These studies report increase in ejection fraction and contractibility, increase in end systolic and diastolic volumes and improvement in the New York Heart Association Class [8, 10, 12, 31]. Although, most authors report a therapeutical benefit this vary enormously among the different studies. This huge variability may reflect differences in number of cells administrated, time between MI and cell therapy, system and site of administration (peripheral or intracoronary infusion, intramyocardial transepicardial or transendocardial injection), criteria for patient selection, repetition of treatment, existence of cardiac surgery at the time of administration, etc.

Clinical trials for cardiac cell therapy are expensive and it takes a long time to obtain reliable data because in most cases differences observed are small and obtaining statistical differences between placebo and treatment requires a large population (which is difficult to obtain when working with human subjects) and, as mentioned above, many parameters need to be considered. Although, at least in part, some of these problems might be overcome with studies in animals, the difference in cardiac physiology between humans and rodents do not guarantee that results can be extrapolated. Therefore, a rational design of clinical trials based on the knowledge of BM-hMSCs effects in the heart is needed if we want to improve the therapeutical efficacy of BM-hMSCs in cardiac cell therapy. A better understanding of how the benefit observed in previous studies is obtained would allow to specifically re-design cellular therapies with BM-hMSCs to improve the outcome in the future.

Different mechanisms have been proposed to explain the capacity of BM-hMSCs to facilitate myocardial repair (review by Psaltis et al. [14]) including: differentiation or transdifferentiation to the cardiac lineage [4, 32], protection of cardiomyocytes against ischemia [4, 15], remodeling of scar tissue [20], activation of the regenerative capacity of resident stem cells [16], modulation of the imflamatory response associated with MI [17, 18], and endothelial differentiation and revascularization of injured area [20-22].

In this study we used in an *in vitro* model of MI that uses cardiomyocytes isolated from neonates rats exposed to an anoxic environment to mimic ischemia (Figure 3). If cells are cultured in glucose free media, the model mimics myocardial ischemia after coronary occlusion when cells are completely deprived of oxygen and die because necrosis in a few hours, but if cultured in glucose containing media cell death takes longer because anaerobic metabolism maintains cell alive in a lethargic/non-contractile state, which probably reflects the situation at the peri-infarcted area. This non-contractile state, which is induced after anoxia-mediated acidification, is reversible if normal culture conditions are restored before the onset of cell death. Therefore, this is a versatile system where is straight forward to test different hypothesis and it is also possible to study the underlying molecular mechanisms.

First, we tested the vasculogenic potential of the BM-hMSCs used in clinical trials for cardiac cell therapy at Hospital Rio Hortega, Valladolid (San-Román et al., manuscript in preparation). These cells, in agreement with the recommendations issued by the International Society for Cellular Therapy [25] expressed CD90, CD105, CD166 but lack the expression of CD34, CD45, HLA-DR, CD117 and CD133, and differentiated to osteoblast when cultured in osteogenic media and to adipoblast when cultured in adipogenic media (Figure 1). In addition, these cells spontaneously differentiated to endothelial cells suggesting that they posses a vasculogenic potential. Several authors have raised the possibility that different subpopulations might account for the multipotency of BM-hMSCs, but the cytometry and clonogenic analysis (Figure 2) indicate that BM-hMSCs used in this study are a fairly homogeneous population of cells that have the potential to differentiate in the three lineages mentioned above.

Previously, Oswald et al. and Al Khaldi et al. [6] described that BM-hMSCs can be differentiated into endothelial cells after treatment with recombinant VEGF [6], a cytokine that plays a fundamental role during formation of new blood vessels and that usually is expressed in response to low O_2_ levels [33]. Interestingly, VEGF was detected at low levels in conditioned media from NRVCs and expression increased more than 10 times when cells were exposed to anoxia (Figure 4A - C). Our data, also confirms that VEGF stimulates the endothelial differentiation of BM-hMSCs (measured as vWF expression) and shows that conditioned media from cardiomyocytes cultured in normoxia or in anoxia also enhances this capacity (Figure 4D-H). Tube formation after culture in matrigel (a measure of the vasculogenic potential of BM-hMSCs) and cell migration across a microporous membrane (cell migration is essential during capillary network development) were also increased by VEGF or conditioned media from NRVCs cultured in anoxia, (Fifure 4I-M, N). Furthermore, VEGF receptor KDR is expressed by BM-hMSCs and downstream mediators of VEGF signaling Akt and p38 are activated after incubation with conditioned media from NRVCs exposed to anoxia. Although, we cannot exclude that other factors may be secreted by NRVCs, we hypothesized that the release of VEGF by NRVCs promotes the vascular differentiation of BM-hMSCs, which, in the context of myocardial infarction, may help to re-vascularize the peri-infarcted region and swing the balance towards recovery of this area by reducing ischemia and thus increasing local contractibility. In fact, when VEGF was blocked in culture supernatants by an antibody anti-VEGF (Figure 5G-J) the capacity of BM-hMSCs to form tubes on matrigel reverted to basal level, indicating that VEGF mediates the pro-angiogenic capacity present in supernatants from NRVCs. In addition, we detected KDR expression and VEGF binding in BM-hMSCs (Figure 5A-D) and VEGF signaling (phosphorylation of Akt and p38) was activated after stimulation with conditioned media from NRVCs cultured in anoxia (Figure 5E).

The low levels of VEGF present in the media during the 7 days that take place the differentiation process may explain the expression of vWF in BM-hMSCs cultured in media from normoxic cardiomyocytes. Surprisingly, expression of late endothelial marker CD31 was not detected suggesting that endothelial differentiation of BM-hMSCs was not complete and suggesting that signals different from VEGF are involved in the maduration.

During the few following days after a heart attack, surrounding the zone of infarction where cells are necrotic, there is an area of weakly or no contracting cells that suffer from mild to moderate ischemia called peri-infarcted area where VEGF expression has been detected (local production of lactate and acidification that occurs during ischemia probably inhibits contractibility in this area [34]). With time, cells within this area die because extended ischemia (during this period rest is recommended because any extra work load in the heart may aggravate the ischemia in this area and accelerate the cell death) or recover contractibility after enlargement of collateral blood vessels and restoration of blood supply, probably stimulated by this VEGF. Finally, necrotic tissue is replaced with scar tissue and the remaining heart tries to compensate for missing musculature, which in case of large scar areas leads to cardiac hypertrophy and congestive heart failure. Therefore, delivery of BM-hMSCs during the recovery period, specially in the ischemic peri-infarcted zone where VEGF is being expressed, may contribute to formation of new capillaries thank to the capacity of these cells to differentiate into the endothelial lineage when stimulated by VEGF. In addition, BM-hMSCs also secrete VEGF in response to anoxia (Figure 5F), which it may act on local vasculature and enhance local angiogenesis.

This would help to reduce the ischemia in the area and therefore would prevent death in this numb tissue, thus reducing the final size of the necrosis. The high resistance to anoxia of BM-hMSCs (Supplemental Figure 1) probably reflects the capacity of these cells to function in ischemic zones.

Consistent with this, in animal models using rodents (both rats and mice), capillary density and myocardial blood flow increased in the infarcted zone, but not in healthy myocardium after intraventricular injection of BM-MSCs, and correlated with an overall improvement in cardiac function. In addition transplanted cells have been identified within new capillaries [29, 35]. Furthermore, similar results have been obtained using CD34+ endothelial progenitor instead of mesenchymal cells [22] confirming a neat benefit from revascularization during cardiac cell therapy. Also, non-cell-based therapeutic angiogenic treatments such as protein-based or gene therapies show promising results [36].

This suggest that in general the benefit of cellular therapies would be higher if cells are delivered directly in the ischemic area needed of revascularization (where circulation is poor) than if cells are delivered through the circulatory system (from where the access to the ischemic area it is more limited). In fact, a comparison between the efficacy of transendocardial injection trough a Biosense NOGA and intracoronary infusion of BM-MSCs in a canine model shows that transendorcardial injection it is by far better: The increase in ejection fraction and in vascularity, and the decrease in myocardial ischemia and end systolic and diastolic volumes are higher with the NOGA system [13]. Consistent with this, a study performed by Villa y co-investigators [37] shows that patients with persistent microvascular obstruction (PMO) fail to show signs of improvement after intracoronary delivery. On the other hand, patients without PMO (in whom cells can get close to the capillary trough coronary delivery) respond as expected.

We also tested the capacity of BM-hMSCs to protect cardiomyocytes against cell death, but we did not find any sign of protection when cardiomyocytes were exposed to anoxia either in glucose free media or in glucose containing media. This contrast from previous studies [15, 29] that have shown that co-culture with BM-hMSCs protects cardiomyocytes against ischemia through Akt activation by VEGF. These differences may be explained by different timing or by exposure to a milder ischemia (1% O _2_ in Dai′s work vs. <0.7% O_2_ in this study as indicated in the technical specifications of BD GasPak pouches). In addition, we found that BM-hMSCs are more resistant to anoxia than NRVCs (Supplemental Figure 1A-B), which may explain, at least in part, previously published data where BM-hMSCs are not identified before determining cell death in co-cultures. However, VEGF expression in ischemic myocardium may serve during physiologic ischemia (as produced during exercise) as a short-term protection while at the same time promotes new vessel formation in the area. In fact, it is well known that periodical exercise increased maximum O_2_ consumption and increases capillary proliferation in the heart [38] and exercise has proved to be beneficial during rehabilitation of patients with coronary heart diseases [39].

Finally, BM-MSCs have the capacity to differentiate in cardiomyocytes [4] and it has been proposed that differentiation of BM-hMSCs into the cardiac lineage may contribute to regenerate the heart muscle after an MI. When we analyzed the expression cardiac MHCαin co-cultures of BM-hMSCs and NRVCs by RT-PCR we did not detect the expression of human MHCα and only MHCα of rat origin was detected after 1 week (Supplemental Figure 1C-D). In addition, when BM-hMSCs were loaded with the cell tracker CMTPX and co-cultured with NRVCs, we did not find expression of MHCαin cells carrying the tracker (Supplemental Figure 1E). Although cellular differentiation it is a slow process and we cannot exclude that differentiation of BM-hMSCs in cardiomyocytes may take longer, these results are in agreement with the modest evidence for cardiac differentiation observed in cardiac cell therapy with BM-hMSCs to date.

## Acknowledgements

This work was supported by Junta de Caslilla y León y Instituto de Salud Carlos III grant PI060480 and. VMC gratefully thanks Dr. Andres Alonso and Dra. María Luisa Nieto for providing us the antibodies used in Western blots.

## References

[1] Haynesworth SE, Goshima J, Goldberg VM, Caplan AI. Characterization of cells with osteogenic potential from human marrow. Bone. 1992; 13(1): 81–8.

[2] Pittenger MF, Mackay AM, Beck SC, Jaiswal RK, Douglas R, Mosca JD, et al. Multilineage potential of adult human mesenchymal stem cells. Science. 1999 Apr 2; 284(5411): 143–7.

[3] Makino S, Fukuda K, Miyoshi S, Konishi F, Kodama H, Pan J, et al. Cardiomyocytes can be generated from marrow stromal cells in vitro. J Clin Invest. 1999 Mar; 103(5): 697–705.

[4] Xu M, Wani M, Dai YS, Wang J, Yan M, Ayub A, et al. Differentiation of bone marrow stromal cells into the cardiac phenotype requires intercellular communication with myocytes. Circulation. 2004 Oct 26; 110(17): 2658–65.

[5] Bruder SP, Jaiswal N, Haynesworth SE. Growth kinetics, self-renewal, and the osteogenic potential of purified human mesenchymal stem cells during extensive subcultivation and following cryopreservation. J Cell Biochem. 1997 Feb; 64(2): 278–94.

[6] Oswald J, Boxberger S, Jorgensen B, Feldmann S, Ehninger G, Bornhauser M, et al. Mesenchymal stem cells can be differentiated into endothelial cells in vitro. Stem Cells. 2004; 22(3): 377–84.

[7] Amado LC, Saliaris AP, Schuleri KH, St John M, Xie JS, Cattaneo S, et al. Cardiac repair with intramyocardial injection of allogeneic mesenchymal stem cells after myocardial infarction. ProcNatlAcadSci U S A. 2005 Aug 9; 102(32): 11474–9.

[8] Chen S, Liu Z, Tian N, Zhang J, Yei F, Duan B, et al. Intracoronary transplantation of autologous bone marrow mesenchymal stem cells for ischemic cardiomyopathy due to isolated chronic occluded left anterior descending artery. J Invasive Cardiol. 2006 Nov; 18(11): 552–6.

[9] Gojo S, Gojo N, Takeda Y, Mori T, Abe H, Kyo S, et al. In vivo cardiovasculogenesis by direct injection of isolated adult mesenchymal stem cells. Exp Cell Res. 2003 Aug 1; 288(1): 51–9.

[10] Katritsis DG, Sotiropoulou PA, Karvouni E, Karabinos I, Korovesis S, Perez SA, et al. Transcoronary transplantation of autologous mesenchymal stem cells and endothelial progenitors into infarcted human myocardium. Catheter Cardiovasc Interv. 2005 Jul; 65(3): 321–9.

[11] Makkar RR, Price MJ, Lill M, Frantzen M, Takizawa K, Kleisli T, et al. Intramyocardial injection of allogenic bone marrow-derived mesenchymal stem cells without immunosuppression preserves cardiac function in a porcine model of myocardial infarction. J CardiovascPharmacolTher. 2005 Dec; 10(4): 225–33.

[12] Mohyeddin-Bonab M, Mohamad-Hassani MR, Alimoghaddam K, Sanatkar M, Gasemi M, Mirkhani H, et al. Autologous in vitro expanded mesenchymal stem cell therapy for human old myocardial infarction. Arch Iran Med. 2007 Oct; 10(4): 467–73.

[13] Perin EC, Silva GV, Assad JA, Vela D, Buja LM, Sousa AL, et al. Comparison of intracoronary and transendocardial delivery of allogeneic mesenchymal cells in a canine model of acute myocardial infarction. J Mol Cell Cardiol. 2008 Mar; 44(3): 486–95.

[14] Psaltis PJ, Zannettino AC, Worthley SG, Gronthos S. Concise review: mesenchymal stromal cells: potential for cardiovascular repair. Stem Cells. 2008 Sep; 26(9): 2201–10.

[15] Dai Y, Xu M, Wang Y, Pasha Z, Li T, Ashraf M. HIF-1alpha induced-VEGF overexpression in bone marrow stem cells protects cardiomyocytes against ischemia. J Mol Cell Cardiol. 2007 Jun; 42(6): 1036–44.

[16] Yoo SW, Kim SS, Lee SY, Lee HS, Kim HS, Lee YD, et al. Mesenchymal stem cells promote proliferation of endogenous neural stem cells and survival of newborn cells in a rat stroke model. Exp Mol Med. 2008 Aug 31; 40(4): 387–97.

[17] Semedo P, Palasio CG, Oliveira CD, Feitoza CQ, Goncalves GM, Cenedeze MA, et al. Early modulation of inflammation by mesenchymal stem cell after acute kidney injury. Int Immunopharmacol. 2009 Jan 13.

[18] Ye Z, Wang Y, Xie HY, Zheng SS. Immunosuppressive effects of rat mesenchymal stem cells: involvement of CD4+CD25+ regulatory T cells. Hepatobiliary Pancreat Dis Int. 2008 Dec; 7(6): 608–14.

[19] Berry MF, Engler AJ, Woo YJ, Pirolli TJ, Bish LT, Jayasankar V, et al. Mesenchymal stem cell injection after myocardial infarction improves myocardial compliance. Am J Physiol Heart Circ Physiol. 2006 Jun; 290(6): H2196–203.

[20] Hou M, Yang KM, Zhang H, Zhu WQ, Duan FJ, Wang H, et al. Transplantation of mesenchymal stem cells from human bone marrow improves damaged heart function in rats. Int J Cardiol. 2007 Feb 7; 115(2): 220–8.

[21] Davani S, Marandin A, Mersin N, Royer B, Kantelip B, Herve P, et al. Mesenchymal progenitor cells differentiate into an endothelial phenotype, enhance vascular density, and improve heart function in a rat cellular cardiomyoplasty model. Circulation. 2003 Sep 9; 108 Suppl 1: II253–8.

[22] Kocher AA, Schuster MD, Szabolcs MJ, Takuma S, Burkhoff D, Wang J, et al. Neovascularization of ischemic myocardium by human bone-marrow-derived angioblasts prevents cardiomyocyte apoptosis, reduces remodeling and improves cardiac function. Nat Med. 2001 Apr; 7(4): 430–6.

[23] Conget PA, Minguell JJ. Phenotypical and functional properties of human bone marrow mesenchymal progenitor cells. J Cell Physiol. 1999 Oct; 181(1): 67–73.

[24] Campa VM, Gutierrez-Lanza R, Cerignoli F, Diaz-Trelles R, Nelson B, Tsuji T, et al. Notch activates cell cycle reentry and progression in quiescent cardiomyocytes. J Cell Biol. 2008 Oct 6; 183(1): 129–41.

[25] Dominici M, Le Blanc K, Mueller I, Slaper-Cortenbach I, Marini F, Krause D, et al. Minimal criteria for defining multipotent mesenchymal stromal cells. The International Society for Cellular Therapy position statement. Cytotherapy. 2006; 8(4): 315–7.

[26] Matsuoka R, Ogawa K, Yaoita H, Naganuma W, Maehara K, Maruyama Y. Characteristics of death of neonatal rat cardiomyocytes following hypoxia or hypoxia-reoxygenation: the association of apoptosis and cell membrane disintegrity. Heart Vessels. 2002 Sep; 16(6): 241–8.

[27] Pugh CW, Ratcliffe PJ. Regulation of angiogenesis by hypoxia: role of the HIF system. Nat Med. 2003 Jun; 9(6): 677–84.

[28] Siekmann AF, Covassin L, Lawson ND. Modulation of VEGF signalling output by the Notch pathway. Bioessays. 2008 Apr; 30(4): 303–13.

[29] Uemura R, Xu M, Ahmad N, Ashraf M. Bone marrow stem cells prevent left ventricular remodeling of ischemic heart through paracrine signaling. Circ Res. 2006 Jun 9; 98(11): 1414–21.

[30] Song YH, Gehmert S, Sadat S, Pinkernell K, Bai X, Matthias N, et al. VEGF is critical for spontaneous differentiation of stem cells into cardiomyocytes. BiochemBiophys Res Commun. 2007 Mar 23; 354(4): 999–1003.

[31] Wang JA, Xie XJ, He H, Sun Y, Jiang J, Luo RH, et al. [A prospective, randomized, controlled trial of autologous mesenchymal stem cells transplantation for dilated cardiomyopathy].ZhonghuaXinXue Guan Bing ZaZhi. 2006 Feb; 34(2): 107–10.

[32] Sanchez A, Fernandez ME, Rodriguez A, Fernandez J, Torre-Perez N, Hurle JM, et al. Experimental models for cardiac regeneration. Nat ClinPractCardiovasc Med. 2006 Mar; 3 Suppl 1: S29–32.

[33] Ferrara N. Molecular and biological properties of vascular endothelial growth factor. J Mol Med. 1999 Jul; 77(7): 527–43.

[34] Samaja M, Allibardi S, Milano G, Neri G, Grassi B, Gladden LB, et al. Differential depression of myocardial function and metabolism by lactate and H+. Am J Physiol. 1999 Jan; 276(1 Pt 2): H3–8.

[35] Tang YL, Zhao Q, Zhang YC, Cheng L, Liu M, Shi J, et al. Autologous mesenchymal stem cell transplantation induce VEGF and neovascularization in ischemic myocardium. Regul Pept. 2004 Jan 15; 117(1): 3–10.

[36] Rosinberg A, Khan TA, Sellke FW, Laham RJ. Therapeutic angiogenesis for myocardial ischemia. Expert Rev Cardiovasc Ther. 2004 Mar; 2(2): 271–83.

[37] Villa A, Tejedor-VinuelaP, Sanchez PL, Tapia C, Arnold R, Gomez-Salvador I, et al. [Effect of persistent microvascular obstruction on post-infarction ventricular remodeling following intracoronary bone-marrow cell transplantation: a contrast-enhanced cardiac magnetic resonance study]. Rev Esp Cardiol. 2008 Jun; 61(6): 602–10.

[38] Anversa P, Levicky V, Beghi C, McDonald SL, Kikkawa Y. Morphometry of exercise-induced right ventricular hypertrophy in the rat. Circ Res. 1983 Jan; 52(1): 57–64.

[39] Taylor RS, Brown A, Ebrahim S,Jolliffe J, Noorani H, Rees K, et al. Exercise-based rehabilitation for patients with coronary heart disease: systematic review and meta-analysis of randomized controlled trials. Am J Med. 2004 May 15; 116(10): 682–92.

